# Role of cytoneme-like structures and extracellular vesicles in *Trichomonas vaginalis* parasite: parasite communication

**DOI:** 10.1101/2023.01.18.524507

**Authors:** Nehuén Salas, Manuela Blasco Pedreros, Tuanne dos Santos Melo, Vanina G. Maguire, Jihui Sha, James A. Wohlschlegel, Antonio Pereira-Neves, Natalia de Miguel

## Abstract

*Trichomonas vaginalis*, the etiologic agent of the most common non-viral sexually transmitted infection worldwide, colonizes the human urogenital tract where it remains extracellular and adheres to epithelial cells. With an estimated prevalence of 276 million new cases annually, mixed infections with different parasite strains are expected. Although it is considered as obvious that parasites interact with their host to enhance their own survival and transmission, evidence of mixed infection call into question the extent to which unicellular parasites communicate with each other. Here, we demonstrated that different *T. vaginalis* strains are able to communicate through the formation of cytoneme-like membranous cell connections. We showed that *T. vaginalis* adherent strains form abundant membrane protrusions and cytonemes formation of an adherent parasite strain (CDC1132) is affected in the presence of a different strain (G3 or B7RC2). Using a cell culture inserts assays, we demonstrated that the effect in cytoneme formation is contact independent and that extracellular vesicles (EVs) are responsible, at least in part, of the communication among strains. In this sense, we found that EVs isolated from G3, B7RC2 and CDC1132 strains contain a highly distinct repertoire of proteins, some of them involved in signaling and communication, among other functions. Finally, we showed that parasite adherence to host cells is affected by this communication between strains as binding of adherent *T. vaginalis* CDC1132 strain to prostate cells is significantly higher in the presence of G3 or B7RC2 strains. Demonstrating that interaction of isolates with distinct phenotypic characteristics may have significant clinical repercussions, we also observed that a poorly adherent parasite strain (G3) adheres more strongly to prostate cells in the presence of an adherent strain. The study of signaling, sensing and cell communication in parasitic organisms will surely enhance our understanding of the basic biological characteristics of parasites that might have important consequences in pathogenesis.

## Introduction

The flagellated protozoan parasite *Trichomonas vaginalis* is the etiologic agent of trichomoniasis, the most common non-viral sexually transmitted infection worldwide with an estimated 276 million new cases annually (WHO, 2018). Although asymptomatic infection is common, multiple symptoms and pathologies can arise in both men and women, including vaginitis, urethritis, prostatitis, low birth weight infants and preterm delivery, premature rupture of membranes, and infertility (Fichorova, 2009; Swygard et al., 2004). *T. vaginalis* has also emerged as an important cofactor in amplifying human immunodeficiency virus (HIV) spread as individuals infected with *T. vaginalis* have a significantly increased incidence of HIV transmission (McClelland et al., 2007; Van Der Pol et al., 2008). In addition, *T. vaginalis* infection increases the risk of cervical and aggressive prostate cancer (Gander et al., 2009; Twu et al., 2014). Due to its great prevalence in some communities, mixed infections with several parasite strains are anticipated. In this sense, an analysis performed of 211 *T. vaginalis* samples isolated in five different continents identified 23 cases of mixed infections (10.9 %) (Conrad et al., 2012). In mixed infections, the extent to which parasites communicate with each other has been severely underestimated.

The fundamental ability to sense, process, and respond to extracellular signals is shared by all living forms. However, little is known about sensing and signaling mechanisms in protozoan parasites compared to other organisms (Roditi, 2016). Although it is widely accepted that pathogens interact with their host to enhance their own survival and transmission, communication between unicellular parasites has been poorly studied (Roditi, 2016). Although it was originally believed that single-celled microorganisms do not need to cooperate with other members of their own species, in recent years it has become clear that microbes are social organisms, capable of communicating with one another and engaging in cooperative behavior (Oberholzer et al., 2010; Roditi, 2016). In this sense, social interactions in a population offer advantages over a unicellular lifestyle that include increased protection from host defenses, access to nutrients, exchange of genetic information and enhanced ability to colonize, differentiate and migrate as a group (Oberholzer et al., 2010; Roditi, 2016).

Cells communicate over short or long distances in different ways. Extracellular vesicles (EVs), soluble secreted factors, membrane protrusions and direct contact between cells are all different forms of cell communication (Buszczak et al., 2016; Matthews, 2021; Regev-Rudzki et al., 2013). Plasma membrane serve as the primary interface between a cell and its environment, mediating direct contact, sensing environmental factors and releasing signaling molecules. Cellular protrusions have emerged as a way for cells to communicate with one another. Among different types of cellular protrusions, filopodia are thin cellular extensions that have been observed in many cell types and have been assigned different roles like cell migration, cell adhesion, force generation, wound healing, environmental sensing, antigen presentation, and neuronal pathfinding (Roy and Kornberg, 2015). Although their physical properties vary (2–400 µm in length, 0.1–0.3 µm diameter), all are actin-based, they extend and retract at velocities that have been measured as much as 25 µm/minute, and their tips can contact other cells (Roy and Kornberg, 2015). Their different shapes and roles are reflected in the many names that have been coined: thin filopodia (Miller et al., 1995), thick filopodia (McClay, 1999), invadopodia (Chen, 1989), telopodes (Popescu and Faussone-Pellegrini, 2010), tunneling nanotubes (Rustom et al., 2004) and cytonemes (Ramírez-Weber and Kornberg, 1999). Specifically, cytonemes are considered as thin specialized filopodia that have been shown to traffic signaling proteins such as morphogens, growth factors and cell determination factors (Roy and Kornberg, 2015). Although cytonemes have similar diameters to conventional filopodia (typically smaller than 200 nm), they have the potential to extend up to ∼300 nm from the originating cell body and have been observed in both vertebrate and invertebrate systems (Kornberg and Roy, 2014).

Alternatively, EVs are also considered as key mediators in intercellular communication in many types of cells. They are a group of heterogeneous particles formed by a lipid bilayer containing proteins and nucleic acids (Abels and Breakefield, 2016; Yáñez-Mó et al., 2015). As proposed by the International Society for Extracellular Vesicles (ISEV), the term ‘‘extracellular vesicles’’ is referred to all sub-populations of EVs, so it is recommended to use it collectively and universally (Théry et al., 2018). Among the various subtypes of extracellular vesicles, the particles can be defined according to the mode of biogenesis, size, and function into three major categories: (1) exosomes formed due to plasma membrane invagination into multivesicular bodies (MVB) with size ranging from 40 to 100 nm; (2) Microvesicles (MVs), also called shedding vesicles, microparticles or ectosomes, originated from the budding and extrusion of the plasma membrane, with sizes between 50 and 1000 nm and an asymmetric structure; and (3) apoptotic bodies, with greater sizes (up to 5000 nm) originated from cells in the process of programed cell death (Kalra et al., 2012). Different to cell membrane protrusions, EVs modulate short- and long-range events, allowing cells to communicate even at long distances. These particles regulate physiological processes, such as blood coagulation, cell differentiation and inflammation, as well as pathological processes caused cancer, neurological, cardiovascular, and infectious diseases (Kao and Papoutsakis, 2019; Raposo and Stoorvogel, 2013; Yáñez-Mó et al., 2015). EVs are relevant for the communication between pathogens and host cells (Nievas et al., 2020, p. 2021; Sabatke et al., 2021; Torrecilhas et al., 2020). Specifically in *T. vaginalis*, the analysis of EVs, both exosomes and MVs, has become a very exciting field in the study of parasite: host interaction as it has been shown that the formation of EVs increase in the presence of host cells and modulate parasite adherence (Nievas et al., 2020; Y R Nievas et al., 2018a; Olmos-Ortiz et al., 2017; Rai and Johnson, 2019; Twu et al., 2013). Although the role of EVs in *T. vaginalis*: host interaction has been deeply analyzed (Nievas et al., 2020; Rada et al., 2022; Rai and Johnson, 2019; Salas et al., 2021; Twu et al., 2013), the understanding of the role of EVs in communication between different parasite strains, and the implications in the infection process, is still scarce.

Here, we demonstrated that different *T. vaginalis* strains can communicate through the formation of cytoneme-like membranous cell connections. We observed that cytoneme formation of an adherent parasite strain (CDC1132) is affected in the presence of a different strain (G3 or B7RC2). Additionally, we demonstrated that the effect in cytoneme formation is contact independent and that extracellular vesicles (EVs) are responsible, at least in part, of the communication between strains. To explain the differential response in cytoneme formation due to the presence of EVs isolated from different strains, we analyzed the EVs protein content by mass spectrometry and demonstrated that highly specific protein cargo was detected in EVs isolated from different strains. Finally, we showed that parasite adherence to host cells is affected by this communication as binding of adherent *T. vaginalis* CDC1132 strain to prostate cells is significantly higher in the presence of G3 or B7RC2 strains. Importantly, we observed that, in the presence of an adherent strain, a poorly adherent parasite strain (G3) adheres more strongly to prostate cells suggesting that interaction between isolates with distinct phenotypic characteristics may have significant clinical repercussions. The study of signaling, sensing and cell communication in parasitic organisms will surely enhance our understanding of the basic biological characteristics of parasites and reveal new potential clinical outcomes.

## Materials and methods

### Parasites, cell cultures and media

*Trichomonas vaginalis* strains G3 (ATCC PRA-98; Beckenham, UK), B7RC2 (ATCC 50167; Greenville, NC, USA) and CDC1132 (Müller et al., 1988) were cultured in TYM (Tryptone-Yeast extract-Maltose) medium supplemented with 10% fetal equine serum and 10 U/ml penicillin (Clark and Diamond, 2002). Parasites were grown at 37°C and passaged daily. The human BPH-1 cells, kindly provided by Dr. Simon Hayward (NorthShore University, USA) (Jiang et al., 2010), were grown in RPMI 1640 medium containing 10% fetal bovine serum (FBS; Internegocios, Argentina) with 10 U/ml penicillin and cultured at 37 °C/5% CO_2_.

### Parasites fluorescent labeling

Parasites were incubated at 37°C for 1 h on glass coverslips as previously described (de Miguel et al., 2010). Parasites attached to coverslips were fixed in 4% paraformaldehyde at room temperature for 20 min and labeled with wheat germ agglutinin (WGA) lectin from *Triticum vulgaris* conjugated with FITC (Sigma). To this end, parasites were incubated 1:100 WGA/phosphate-buffered saline (PBS) dilution at 37°C during 1 h, washed three times with PBS solution and mounted using Fluoromont Aqueous Mounting Medium (Sigma). Fluorescent parasites were visualized using a Zeiss Axio Observer 7 (Zeiss) microscope.

### Scanning Electron Microscopy

Cells were washed with PBS solution and fixed in 2.5% glutaraldehyde in 0.1 M cacodylate buffer, pH 7.2. The cells were then post-fixed for 15 min in 1% OsO_4_, dehydrated in ethanol and critical point-dried with liquid CO_2_. The dried cells were coated with gold–palladium to a thickness of 15 nm and then observed with a Jeol JSM-5600 scanning electron microscope, operating at 15 kV. Some images were colored using Adobe Photoshop software (Adobe USA), version 24.0.1.

### Parasite aggregation

Clumps formation was analyzed in parasites grown in regular TYM media under anaerobic conditions at different concentration. A clump was defined as the size corresponding to an aggrupation of at least 5 parasites. Quantification of clumps in thirty 20X magnification fields was performed using a Nikon TSM (Nikon) microscope. Three independent experiments were performed.

### Direct co-culture of parasites

CDC1132 parasites were labelled using CellTracker Red CMTPX Dye (ThermoFisher). Then, labelled CDC1132 parasites were co-incubated for 1 hour at 37°C with unlabeled G3, B7RC2 and CDC1132 parasites at different cell ratios (1:1, 1:2, and 1:9). Parasites attached to coverslips were fixed and labelled with WGA as described previously. The number of CDC1132 parasites containing filopodia and cytonemes, visualized as the ones with double labelling (red and green), was analyzed using a Zeiss Axio Observer 7 (Zeiss) microscope. Three independent experiments, each of them in duplicates, were performed.

### Indirect parasites co-culture using cell culture inserts assays

Parasites from G3, B7RC2 and CDC1132 strains were co-cultured using polyester transwell-24 inserts (1 µm pore size; Biofil). CDC1132 parasites were loaded at the bottom of each well and exposed to parasites from G3, B7RC2 and CDC1132 strains placed into the transwell inserts (parasite ratio 1:2 bottom: transwell) for 1 hour at 37°C. Then, CDC1132 parasites attached to coverslips were labelled with WGA and the formation of filopodia and cytonemes was analyzed using a Zeiss Axio Observer 7 (Zeiss) microscope.

### Isolation of T. vaginalis extracellular vesicles

As recommended, the term is referred to all sub-populations of EVs including exosome and microvesicles (MVs) (Théry et al., 2018). As previously described (Salas et al., 2021), EVs were isolated in parallel from 250 ml cultured parasites (10^6^ cells/ml) from G3, B7RC2 and CDC1132 strains by incubating the parasites for 4 h at 37°C in TYM medium without serum. Then, conditioned medium was harvested and centrifuged at 3000 rpm for 10 min to remove cell debris. The media was filtered through a 0.8 μm filter and the sample was pelleted by centrifugation at 100,000 x g for 90 min to obtain an EVs enriched fraction (a mixture of MVs and exosomes). The pellet was resuspended in 200 μl cold PBS + 1X cOmplete™ ULTRA Tablets, Mini, EASYpack Protease Inhibitor Cocktail (Sigma). EVs isolation from G3, B7RC2 and CDC1132 were performed in parallel. Three independent experiments were performed.

### Total protein quantification

Total protein concentration was determined colorimetrically (Bradford Reagent, Sigma-Aldrich). The standard curve was prepared using Bovine Serum Albumin (Promega). Absorbance was measured at 595 nm with a spectrophotometer.

### Incubation of parasites with EVs

To carry out the incubation step, 10 and 20 μg of EVs or the same volume of the PBS solution (control) were incubated for 1 hour at 37ºC with CDC1132 parasites in a 24-well plate. Then, parasites were washed twice with PBS and fixed using 4% paraformaldehyde for 20 min. Parasites attached to coverslips were labelled with WGA as previously described and the formation of cytonemes and filopodia was examined using a Zeiss Axio Observer 7 (Zeiss) inverted fluorescence microscope. Three independent experiments were performed in duplicates.

### Proteomic mass spectrometry analysis

EVs enriched samples were resuspended in a minimal volume of digestion buffer (100 mM Tris–HCl, pH 8, 8 M urea). Resuspended proteins were reduced and alkylated by the sequential addition of 5 mM tris(2-carboxyethyl) phosphine and 10 mM iodoacetamide as described previously. The samples were then digested by Lys-C (Princeton Separations) and trypsin proteases (Promega) (Florens et al., 2006). First, Lys-C protease [∼1:50 (w/w) ratio of enzyme: substrate] was added to each sample and incubated for 4 h at 37°C with gentle shaking. The digests were then diluted to 2 M urea by the addition of digestion buffer lacking urea, and trypsin was added to a final enzyme: substrate ratio of 1:20 (w/w) and incubated for 8 h at 37°C with gentle shaking. Digests were stopped by the addition of formic acid to a final concentration of 5%. Supernatants were carefully removed from the resin and analyzed further by proteomics mass spectrometry. Digested samples were then analyzed using a LC–MS/MS platform as described previously (Kaiser and Wohlschlegel, 2005; Wohlschlegel, 2009). Briefly, digested samples were loaded onto a fused silica capillary column with a 5-μm electrospray tip and packed in house with 18 cm of Luna C18 3 μM particles (Phenomenex). The column was then placed in line with a Q-exactive mass spectrometer (ThermoFisher) and peptides were fractionated using a gradient of increasing acetonitrile. Peptides were eluted directly into the mass spectrometer, where MS/MS spectra were collected. The data-dependent spectral acquisition strategy consisted of a repeating cycle of one full MS spectrum (resolution = 70,000) followed by MS/MS of the 12 most intense precursor ions from the full MS scan (resolution = 17,500) (Kelstrup et al., 2012). Raw data and spectra analyses were performed using the MaxQuant software (Tyanova et al., 2016). For protein identification a search against a fasta protein database was done consisting of all predicted open reading frames downloaded from TrichDB on November 9, 2022 (Amos et al., 2022) concatenated to a decoy database in which the amino acid sequence of each entry was reversed. The following searching parameters were used: (1) precursor ion tolerance was 20 ppm; (2) fragment ion tolerance was 20 ppm; (3) cysteine carbamidomethylation was considered as a static modification; (4) peptides must be fully tryptic; and (5) no consideration was made for missed cleavages. False positive rates for peptide identifications were estimated using a decoy database approach and then filtered using the DTASelect algorithm (Cociorva et al., 2006; Elias and Gygi, 2007; Tabb et al., 2002). Proteins identified by at least two fully tryptic unique peptides, each with a false positive rate of less than 5%, were considered to be present in the sample. Three different sets of EVs enriched samples isolated from G3, B7RC2 and CDC1132 strains were independently analyzed. Proteins present in the EVs fraction were identified using BLAST tool (Basic Local Alignment Search Tool) and classified using the GO term enrichment according to PANTHER Classification System (Mi et al., 2013).

### Attachment assay

A modified version of an *in vitro* assay to quantify the binding of *T. vaginalis* to host cell monolayers was carried out (Bastida-Corcuera et al., 2005). Briefly, BPH-1 cells were seeded on coverslips in 24-well plates with RPMI culture medium (Invitrogen) and grown to confluence at 37°C and 5% CO_2_. Parasites (CDC1132 or G3 strain) were labelled with Cell Tracker Blue CMAC (7-amino-4-chloromethylcoumarin) (Invitrogen), added to confluent BPH-1 cells (1:3 parasite: host cell ratio) and exposed to different parasites strains loaded in the inserts in a 1:2 bottom: transwell ratio. Plates were then incubated together at 37°C and 5% CO_2_ for 60 min. Coverslips were subsequently washed in phosphate-buffered saline (PBS) solution, fixed with 4% paraformaldehyde and mounted on slides with Fluoromont Aqueous Mounting Medium (Sigma). Quantification of fluorescent parasites attached to host cells were measured using a Zeiss Axio Observer 7 microscope. Thirty 10X magnification fields were analyzed per coverslip. All experiments were performed three independent times in duplicates.

### Graphics and statistical analyses

Specific statistical considerations and the tests used are described separately for each subsection. GraphPad Prism for Windows version 8.00 was used for graphics. Data are shown as mean ± standard deviation (SD). Statistical significance was established at p< 0.05 and for statistical analyses the InfoStat software (Di Rienzo et al., 2011) version 2020e was used.

## Results

### T. vaginalis adherent strains form abundant membrane protrusions

Visualization of *T. vaginalis* by fluorescent and live microscopy revealed the presence of filamentous structures extending from the surface of some cells (Sup. Video 1). Due to their morphological appearance (Fig. 1A), these structures resembled previously described filopodia and cytonemes (Kornberg and Roy, 2014). To determine if these structures were related to pathogenesis, we evaluated the presence of filopodia and cytonemes in strains with different adherence capacities to host cells: two poorly adherent strains (G3 and NYU209) and two highly adherent strains (B7RC2 and CDC1132) using the membrane binding lectin wheat germ agglutinin (WGA). When the number of membrane protrusions was quantified, we observed that the highly adherent strains B7RC2 and CDC1132 have greater number of filopodia and cytonemes compared to poorly adherent strains G3 and NYU209 (Fig. 1A). In concordance, scanning electron microscopy (SEM) revealed that multiple filopodia and cytonemes originated from the surface of highly adherent CDC1132 parasites and almost no such protrusions were observed in the poorly adherent G3 strain (Fig. 1B). These tubular extensions appear to be close ended and can be seen protruding from the region where the flagella emerge as well as from the cell body, with a length that may reach up to 4.6 µm (Fig. 1B; Suppl. Fig. S1). The cytonemes from the flagellar base are unbranched, displaying a homogeneous morphology and size ranging from 70 nm to 100 nm in thickness (Fig. 1B; Suppl. Fig. S1). Their ultrastructural features are similar to those flagellar projections previously described in the related veterinary trichomonad *Tritrichomonas foetus* (Benchimol et al., 2021). However, the tubular extensions from the cell body are heterogeneous, with diameter varying from 70 nm to 540 nm and some of them furcated (Fig. 1B; Suppl. Figs. S1-S2). Altogether, these findings demonstrate that *T. vaginalis* contains different types of membrane protrusions arising from their surface and suggest that its abundance depends, at least in part, on the phenotype of the strain.

**Figure 1:**
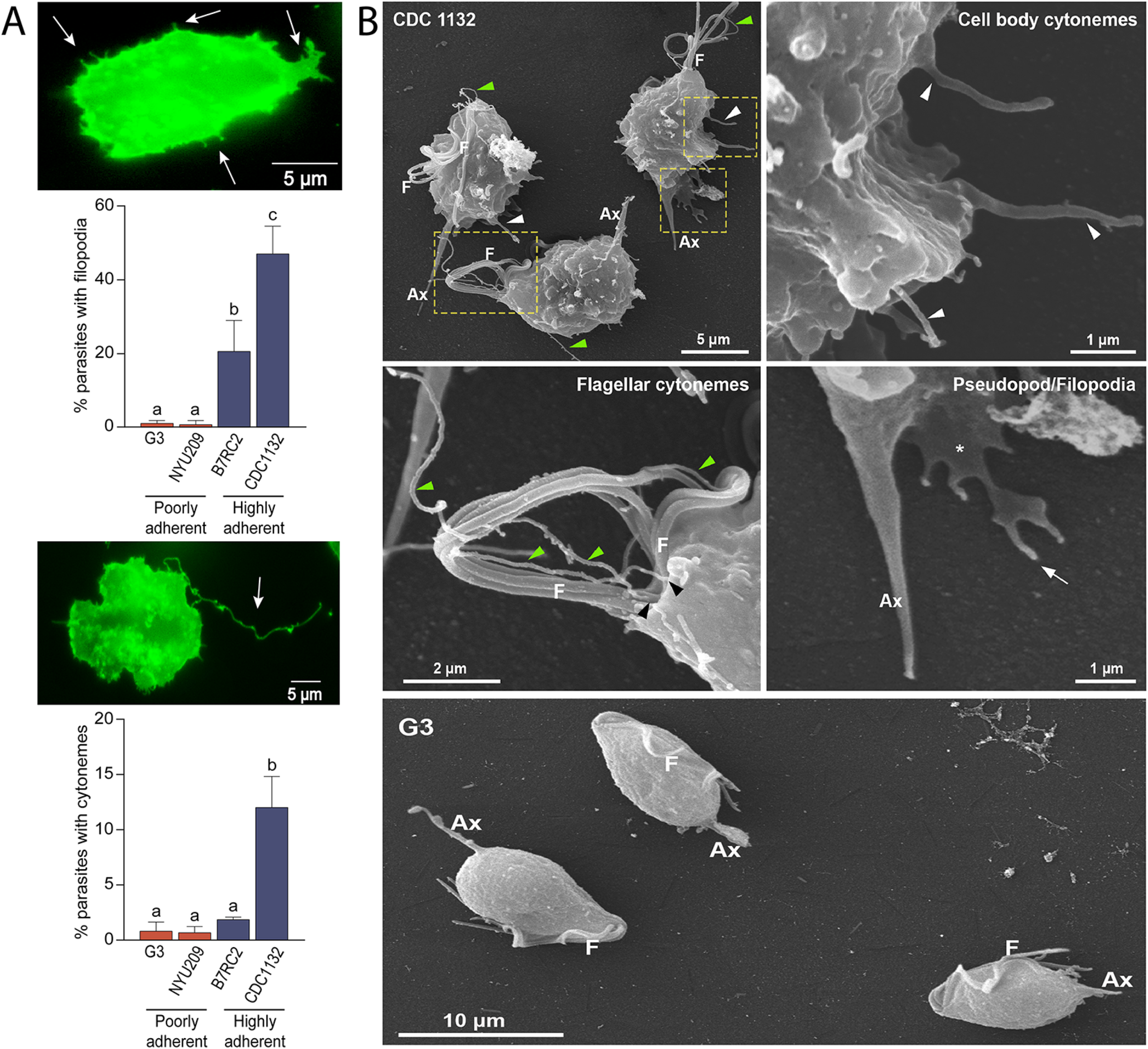
Adherent *T. vaginalis* strains form abundant membrane protrusions. **A**. Quantification of the percentage of parasites containing filopodia (top) or cytonemes (bottom) in their cell surface. The presence of filopodia/cytonemes in two poorly adherent (G3 and NYU209) and two highly adherent strains (B7RC2 and CDC1132) was analyzed. Three independent experiments by duplicate were performed and 100 parasites were randomly counted per sample. Data are expressed as percentage of parasites with filopodia and cytonemes ± standard deviation (SD). ANOVA followed by Tukey’s post hoc test was used to determine significant differences. **B**. Scanning electron microscopy reveals a myriad of projections originating from the surface of parasites from CDC1132 strain. Cytonemes protruding from cell body and flagellar base region are indicated by white and green arrowheads, respectively. Pseudopodia (*) and filopodia (arrow) are also seen. The surface protuberances appear to be close ended. Almost no protrusions were observed arising from the surface of poorly adherent parasites (G3). F, flagella; Ax, axostyle.

### Cytonemes are associated with parasite clumps formation

It has been previously shown that formation of clumps in cell culture generally correlates with the ability of the strain to adhere and be cytotoxic to host cells (Coceres et al., 2015; Lustig et al., 2013). Specifically, highly adherent strains tend to aggregate when cultured in the absence of host cells in contrast to poorly adherent strains that generally do not form clumps *in vitro* (Coceres et al., 2015; Y R Nievas et al., 2018b). Based on this observation and our results by SEM demonstrating that cytonemes are usually detected connecting different parasites inside the clumps (Fig. 2A), we evaluated whether the formation of clumps (Fig. 2B) is accompanied with an increase in number of parasites containing cytonemes (Fig. 2C). As can be observed in figure 2, the number of parasites containing cytonemes is higher at 10^6^ parasites/ml (Fig. 2C); a cell culture condition where usually clumps are formed (Fig. 2B). These results suggest that cytonemes might be involved in parasite: parasite communication. In concordance, cytonemes are usually observed connecting two parasites (Fig. 2D). The closed tip of cytonemes was frequently seen in contact with any region of the surface of an adjacent parasite (Suppl. Fig. S2F), including other cytonemes (Figs 2A, 2D).

**Figure 2:**
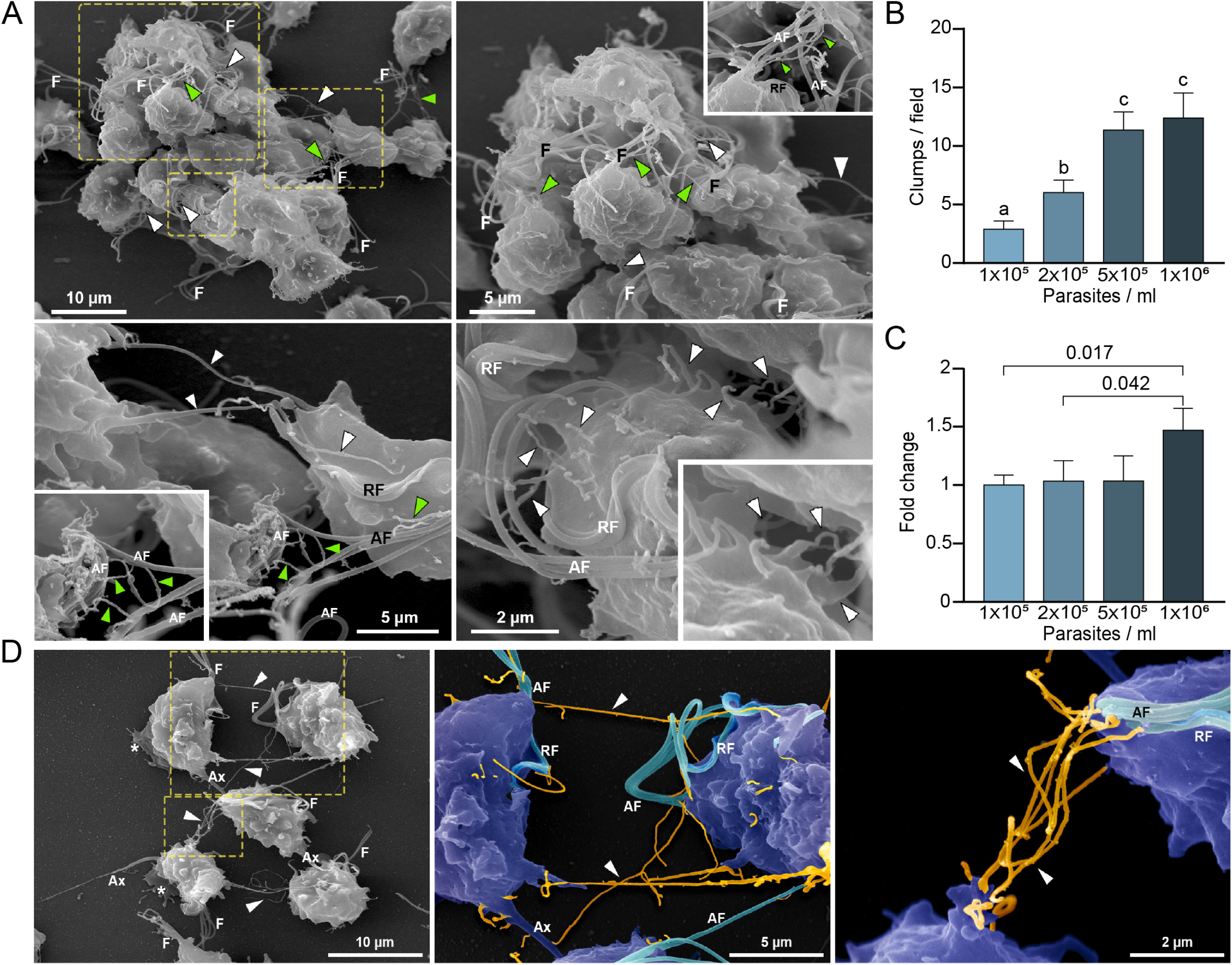
Cytonemes are associated to clumps formation. **A**. Cytonemes from flagella base (green arrowheads) and cell body (white arrowheads) are frequently observed inside clumps of parasites by SEM, connecting several regions of adjacent cells. F, flagella; AF, anterior flagella; RF, recurrent flagellum. **B**. Quantification of clumps per field at different parasite densities (parasites/ml). Twenty fields were counted by duplicate in three independent experiments. A clump was defined as an aggregate of ∼5 or more parasites. ANOVA followed by Tukey’s post hoc test were used to determine significant differences. **C**. Quantification of CDC1132 parasites containing cytonemes at different parasite densities (parasites/ml). Three independent experiments by duplicate were performed and 100 parasites were randomly counted per sample. Student T-tests (α =0.95) were used to determine significant difference between treatments. **D**. Cytonemes (orange) are observed connecting two parasites (blue) by SEM (arrowheads). F, flagella; AF, anterior flagella; RF, recurrent flagellum; Ax, axostyle.

### Interaction between different strains induce formation of membrane protrusions

Due to the great prevalence of *T. vaginalis*, mixed infections with different parasite strains have been observed (Conrad et al., 2012). However, the extent to which parasites communicate and interact with each other during infection has never been analyzed. To evaluate if different parasite strains are able to interact, CDC1132 parasites stained with CFSE (green) were co-incubated for 1 hour with a highly adherent strain (B7RC2) or a poorly adherent strain (G3) stained with CellTracker CMTPX Dye (red). As control, CDC1132 stained with CFSE was also co-incubated with CDC1132 stained with CellTracker CMTPX Dye (red). As can be observed in Figure 3A, CDC1132 parasites are able to interact and form clumps with parasites from B7RC2 or G3 strains (Fig. 3A). As G3 is a poorly adherent strain and usually do not form clumps (Coceres et al., 2015), the formation of smaller clumps when incubated with CDC1132 is expected (Fig. 3A). On the basis of these findings and the observation that cytoneme are usually found connecting parasites (Fig. 2D), we hypothesized that communication among different *T. vaginalis* strains might involve cytonemes and/or filopodia formation. To evaluate this, CDC1132 strain was stained with CellTracker CMTPX Dye (red) and co-incubated with varying amount of unstained G3, B7RC2 and CDC1132 strains (1:1; 1:2; 1:9 ratio). Then, co-incubated parasites were stained with WGA and the formation of filopodia and cytonemes in CDC1132 strain was evaluated by fluorescent microscopy (CDC1132 was visualized as the cells stained with red and green). Although we could not observe a clear increase in filopodia formation in CDC1132 strain upon exposure to G3 or B7RC2, the formation of cytonemes in CDC1132 is clearly affected by the presence of a different strain compared to the exposure to CDC1132 itself (Fig. 3B). These results suggest that cytonemes are specifically involved in parasite: parasite communication.

**Figure 3:**
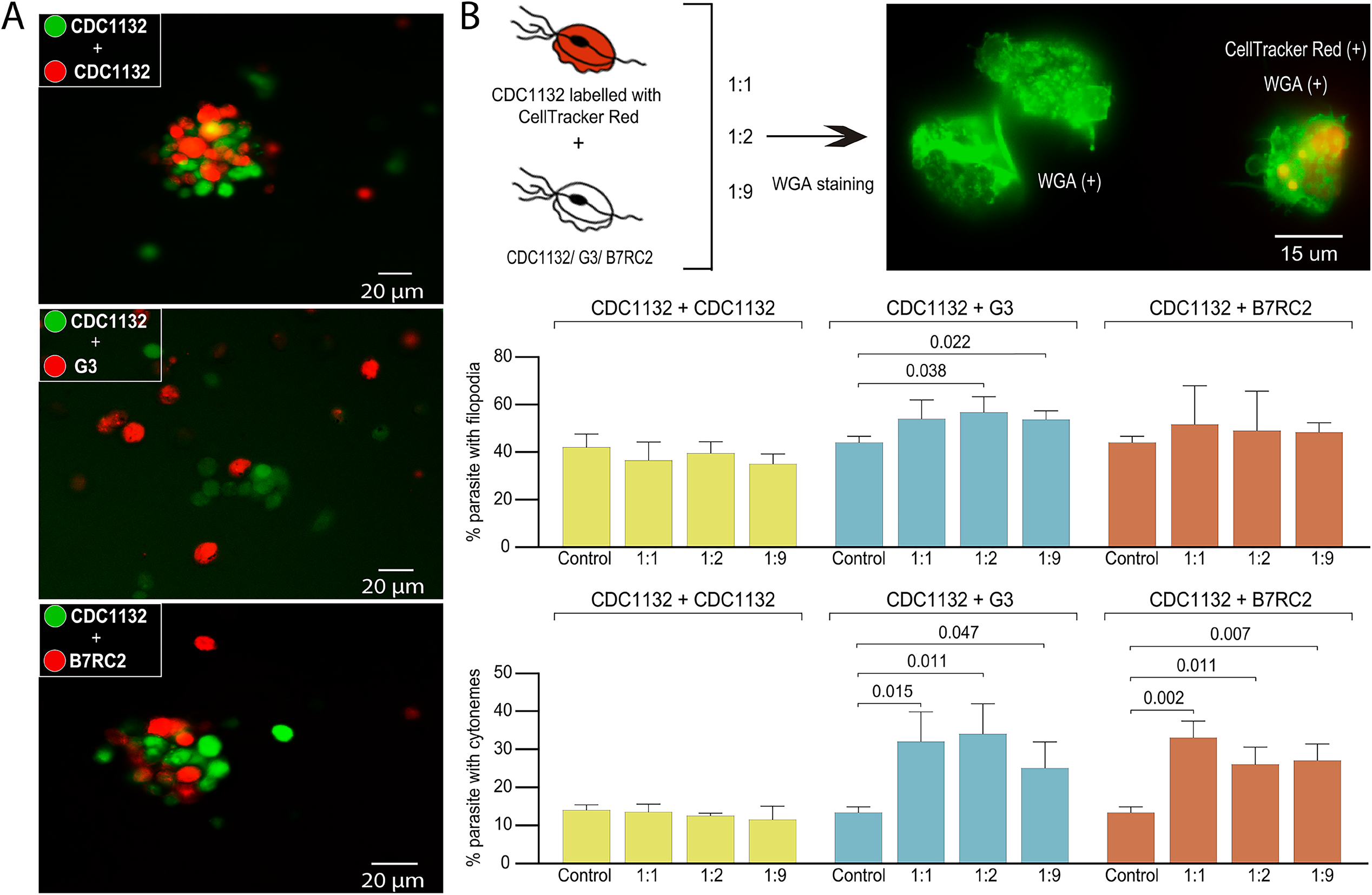
Cytoneme formation is induced by interaction between different strains. **A**. CDC1132 parasites (green) were co-incubated with CDC1132, G3 or B7RC2 (red) for 1 hour. The interaction between different strains was evaluated by analyzing the capacity to form clumps. **B**. Percentage of CDC1132 parasites containing filopodia or cytonemes during co-incubation with different strains at different ratio (1:1, 1:2 and 1:9). CDC1132 parasites were stained with Cell Tracker Red (red) and co-incubated for 1 h with unstained CDC1132, G3 and B7RC2. Then, all the parasites were stained with WGA (green) and the number of CDC1132 parasites containing filopodia and cytoneme (stained red and green) were analyzed. Data are expressed as percentage of parasites with filopodia and cytonemes ± standard deviation (SD). Student T-tests (α =0.95) were used to determine significant difference between treatments.

### Strain specific parasite-secreted extracellular vesicles promote cytoneme formation

To evaluate if the observed increase in cytoneme formation in CDC1132 strain due to the presence of a different strain is the result of physical cell contact, we exposed the CDC1132 strain to G3, B7RC2 and CDC1132 (control) strains using a 1 µm porous membrane that prevent direct contact between parasites (Fig. 4A). This system allows for secreted factors and small extracellular vesicles to pass between the two cell populations but keeps the parasites (10 to 15 µm in diameter) loaded in the inserts from contacting the parasites inoculated in the bottom. To measure paracrine signal activation, the number of CDC1132 parasites in the bottom of the well containing cytonemes and filopodia was measured by WGA staining after 1 h incubation (Fig. 4A). As can be observed in Figure 4A, the number of receiving parasites containing cytonemes and filopodia significantly increase when G3 or B7RC2 strains are loaded in the inserts, compared to the control where CDC1132 strain was loaded in the chamber. These results demonstrated that the effect in cytoneme formation is, at least partially, contact independent. Additionally, these results also suggest that cytoneme formation is affected by paracrine communication among *T. vaginalis* strains.

**Figure 4:**
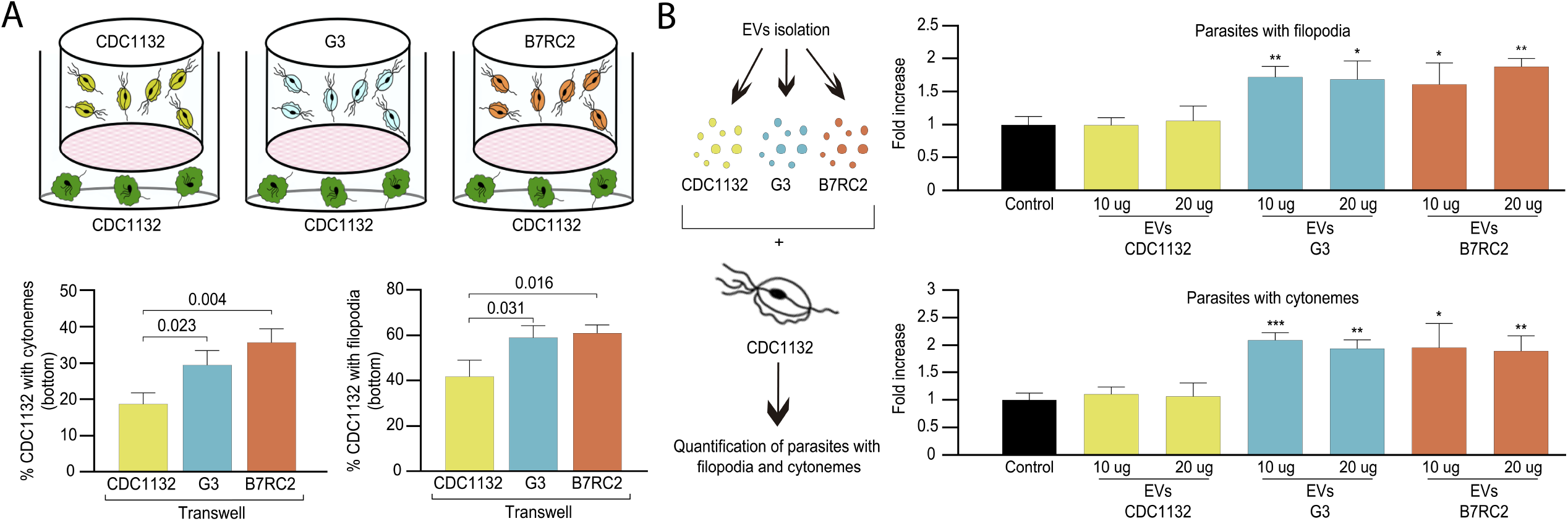
Cytoneme formation is induced by paracrine communication. **A**. Using a cell culture insert assay, CDC1132 parasites (bottom) were co-cultured with CDC1132, G3 and B7RC2 strains (transwell) for 1 h. Then, the number of CDC1132 parasites containing filopodia and cytonemes (bottom) was quantified by WGA staining. Data are expressed as percentage of parasites with filopodia and cytonemes ± standard deviation (SD). Student T-tests (α =0.95) were used to determine significant difference between treatments. **B**. Extracellular vesicles (10 or 20 µg) isolated from G3, B7RC2 and CDC1132 parasites were incubated with wild-type CDC1132 parasites for 1 h. As control, CDC1132 parasites were incubated with the same volume of the PBS solution. Then, the parasites were stained with WGA and the number of parasites containing cytonemes and filopodia was quantified. Data are expressed as a mean -fold increase compared to control (without EVs) ± standard deviation. Student T-tests (α =0.95) were used to determine significant difference between treatments (*p <0.05, **p <0.01, ***p <0.001).

As the cell culture inserts system allows for secreted factors and extracellular vesicles (EVs) to pass throughout the filter, we next asked whether EVs are responsible of the observed cytoneme increased formation during co-incubation with a different parasite strain. Therefore, we isolated EVs from G3, B7RC2 and CDC1132 strains using a previously established protocol (Salas et al., 2021) and enriched EVs samples were incubated directly with CDC1132 parasites (Fig. 4B). Addition of 10 µg EVs from G3 or B7RC2, but not EVs from CDC1132, increased the formation of cytonemes and filopodia of CDC1132 recipient parasites (Fig. 4B). These results suggest that specific molecules are loaded in EVs isolated from different strains.

### Highly specific protein cargo was detected in extracellular vesicles isolated from different strains

To explain the differential response in cytoneme and filopodia formation induced by the presence of EVs isolated from different strains, we analyzed the EVs protein content by mass spectrometry. Three individual samples of EVs isolated from G3, B7RC2 and CDC1132 parasite strains were analyzed (Fig. 5A). Principal-component analysis (PCA) confirmed that the signatures of EVs enriched samples isolated from the different strains were clearly distinct (Fig. 5A). Overall, we identified 1317, 954 and 375 proteins in the EVs enriched samples isolated from G3, CDC1132 and B7RC2 strains, respectively (Fig. 5B and Supplementary Table 1). Although EVs has been isolated from the same number of parasites, the number of proteins identified in the proteome from G3 and CDC1132 was consistently higher than the number of proteins present in B7RC2 samples (Fig. 5B). Comparison of proteins detected in the EVs proteomes from G3, B7RC2 and CDC1132 with previously published *T. vaginalis* secretome (Štáfková et al., 2018), sEVs (Govender et al., 2020; Rada et al., 2022; Twu et al., 2013), EVs (Salas et al., 2021), MVs (Yesica R. Nievas et al., 2018) and/or surface proteome (de Miguel et al., 2010) showed significant overlap as 89% of the proteins identified were previously detected in other proteomes (Table 1S). When the protein content was analyzed, the results indicated that only a core of 346 proteins is shared among EVs isolated from different strains (Fig. 5B) as only. Similarly, the dendrogram that resulted from a hierarchical clustering analysis of proteins using Pearson correlation as a distance metric demonstrated that the proteins of EVs samples from the different strains were definitely distinct from each other (Fig. 5C). As expected, genes involved in several biological processes were highly enriched using the Gene Ontology (GO) analysis (Table 1S and Fig. 5D). Even while the identity of proteins detected in the EVs of each strain differ from one another, the GO analysis indicates that the biological processes detected in the different EVs preparations are highly conserved (Fig. 5D). Specifically, proteins associated to cellular and metabolic processes, response to stimulus, signaling, developmental process and locomotion have been detected in EVs enriched samples isolated from G3, B7RC2 and CDC1132 (Fig. 5D). The diversity of proteins detected in the analyzed EVs proteomes suggest that may serve as part of a strain specific sorting pathway for delivery of biologically active molecules to neighboring cells.

**Figure 5:**
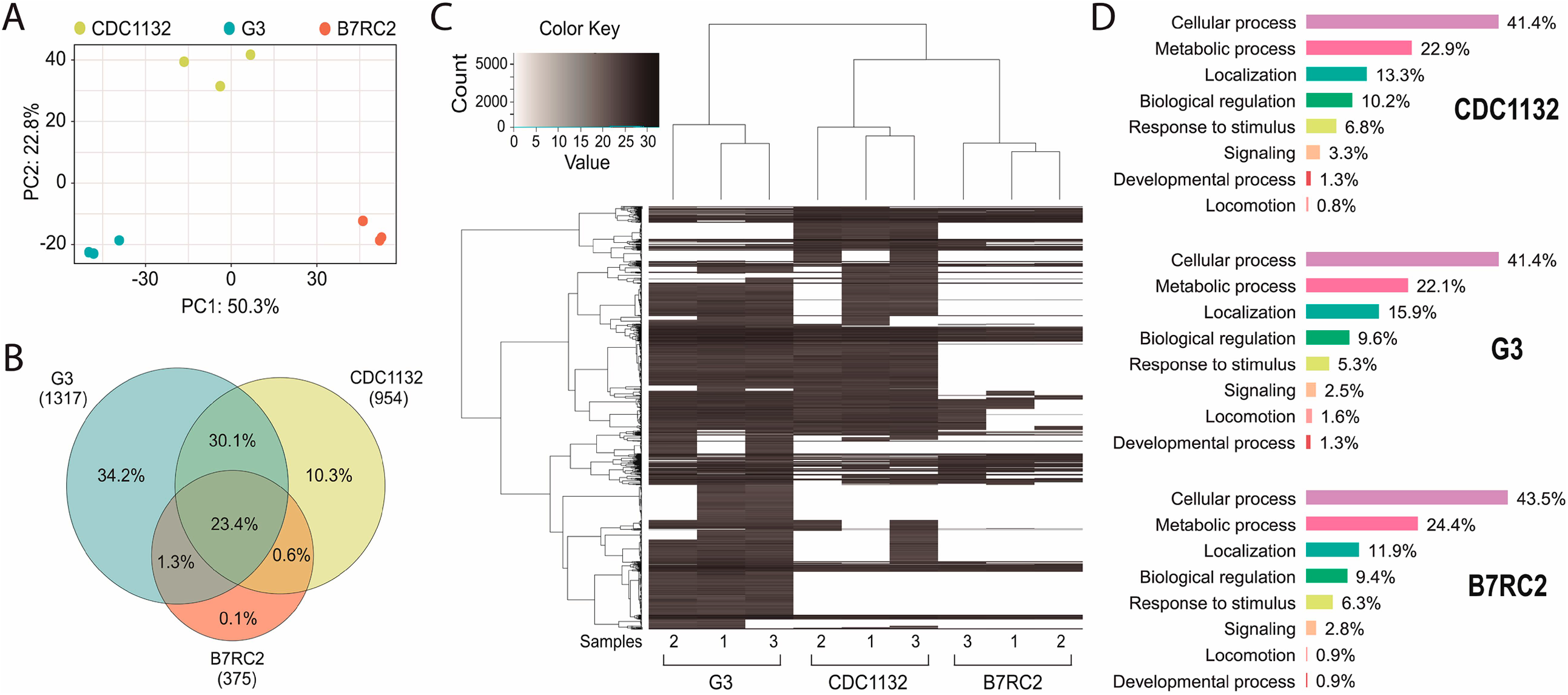
Extracellular vesicles isolated from different strains contain different protein cargo. **A**. Principal component analysis (PCA) plot representing proteomics data from the comparative analysis of 3 independent samples of extracellular vesicles isolated from CDC1132, G3 and B7RC2 strains. **B**. Venn diagram depicting proteins shared among EVs isolated from G3, B7RC2 and CDC1132 strains. **C**. Heatmap of proteins present in EVs of G3, B7RC2 and CDC1132 strains. Each horizontal line representing an individual protein. Color gradient represents the changes of protein abundance. **D**. Proteins were identified using BLAST analysis and sorted into functional groups based on Gene Ontology biological processes.

### Communication between different T. vaginalis strains affect parasite adherence to prostate BPH-1 cells

For extracellular pathogens such as *T. vaginalis*, the ability of a parasite to adhere to its host is likely a determinant of pathogenesis. To attach, *T. vaginalis* changes morphology within minutes: the flagellated free-swimming cell converts into the amoeboid-adherent stage (Kusdian et al., 2013). As it is well-established the connections between pathogenesis and ameboid morphological transitions in *T. vaginalis* (Kusdian et al., 2013) and our results here, we investigated whether the presence of a different parasite strain could affect the ability of the recipient parasite to convert to an ameboid form (Fig. 6A). Using a cell culture inserts assay, we observed that the percentage of amoeboid CDC1132 cells is higher when G3 or B7RC2 are loaded on the inserts compared to the presence of CDC1132 in the chamber (Fig. 6A). These results indicates that communication among different *T. vaginalis* strain induce ameboid transformation in recipient parasites.

**Figure 6:**
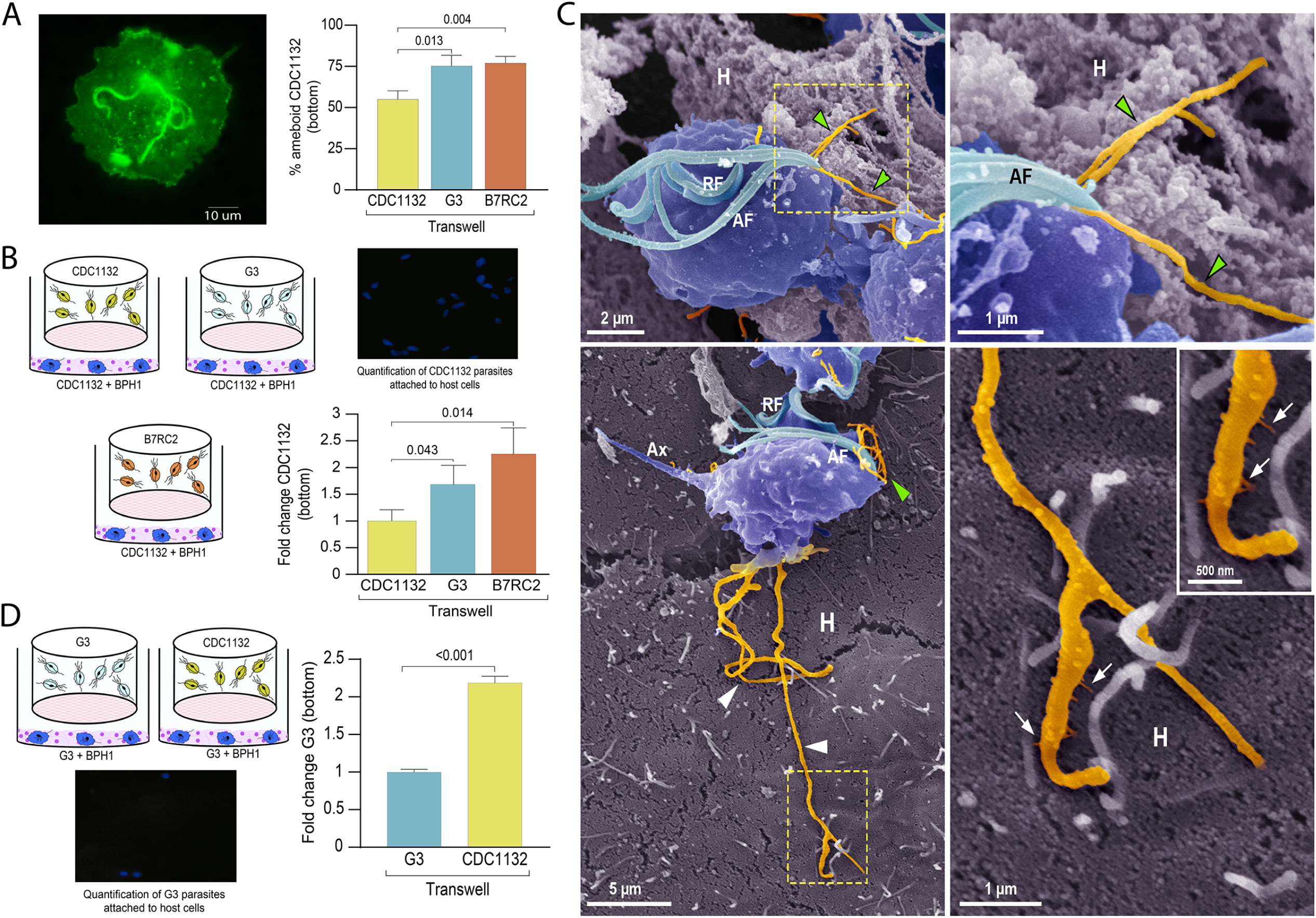
Communication between different parasite strains affects attachment to the host cell. **A**. Using a cell culture insert assay, CDC1132 parasites (bottom) were co-cultured with CDC1132, G3 and B7RC2 strains (transwell) for 1 h. Then, the percentage of amoeboid CDC1132 parasites (bottom) was quantified by WGA staining. Data are expressed as percentage of ameboid parasites ± standard deviation (SD). Student T-tests (α =0.95) were used to determine significant difference between treatments. **B**. Cell Tracker Blue CMAC labelled CDC1132 parasites were incubated for 60 minutes at 37 °C with NhPRE1 prostate cell monolayers cultured onto coverslips in 24-well plates, accompanied by co-culture with CDC1132, G3 and B7RC2 utilizing a cell culture insert assay (1:2 bottom: transwell parasite ratio). Coverslips were washed to remove non-attached parasites, mounted, and quantified by fluorescence microscopy. Data are expressed as fold change related to CDC1132 parasites incubated with CDC1132 strain inside the transwell ± the standard error of the mean (SEM). Student T-tests (α =0.95) were used to determine significant difference between treatments. **C**. Cytonemes (orange) protruding from the flagellar base (green arrowheads) and cell body (white arrowheads) of CDC1132 parasites (blue) are observed in contact with host cells (H) by SEM. In the lower panels, thin extensions branching (arrows) from the cytoneme are seen in close contact with the BPH1 cells. AF, anterior flagella; RF, recurrent flagellum; Ax, axostyle. **D**. Cell Tracker Blue CMAC labeled G3 parasites were incubated for 60 minutes at 37 °C with NhPRE1 prostate cell monolayers cultured on coverslips in 24-well plates, accompanied by co-culture with G3 and CDC1132 utilizing a cell culture insert assay (1:2 bottom: transwell parasite ratio). Coverslips were washed to remove non-attached parasites, mounted, and quantified by fluorescence microscopy. Data are expressed as fold change related to G3 parasites incubated with G3 strain in the transwell ± the standard error of the mean (SEM). Student T-tests (α =0.95) were used to determine significant difference between treatments.

As *T. vaginalis* amoeboid forms have been previously associated to parasite adherence (Arroyo et al., 1993), we then evaluated if communication among strains could affect the adherence of CDC1132 to BPH1 prostate cells. To this end, we performed *in vitro* binding to prostate BPH1 cells experiments using a membrane cell culture inserts system that allow co-incubation of different *T. vaginalis* strains (Fig. 6B). Results demonstrate that binding of *T. vaginalis* CDC1132 strain to BPH1 is significantly higher when G3 or B7RC2 are loaded on the insert as opposed to the presence of CDC1132 in the chamber (Fig. 6B). These results could be suggesting that communication among different *T. vaginalis* strain might affect the behavior of recipient parasites. In concordance with our previous results demonstrating that parasite communication induces cytoneme formation (Fig. 3) and increase in host cell adherence (Fig. 6B), we frequently observed cytonemes protruding from both flagellar base and cell body of CDC1132 in contact with BPH1 cells by SEM (Fig. 6C and Video Sup. 2). Thin extensions branching from the cytoneme were also seen in close contact with the BPH1 cells (Fig. 6C).

To test whether the parasite: parasite communication could affect the behavior of a poorly adherent strain, we expanded our attachment assay and examined whether communication between a highly adherent *T. vaginalis* strain (CDC1132) and a poorly adherent strain (G3) could affect the attachment of G3 to BPH1 cells. Importantly, co-incubation of CDC1132 strain with G3 parasites resulted in a 2-fold increase in G3 attachment to BPH1 (Fig. 6D). In concordance with previous reports (Twu et al., 2013), these data indicate that conditions media from a highly adherent strain can increase parasite attachment to host cells of a less adherent strain. These results suggest that interaction of isolates with distinct phenotypic characteristics may have significant clinical repercussions during mixed infections.

## Discussion

All living organisms, including pathogens, sense extracellular signals and communicate with other cells. Parasites are social organisms, capable of communicating with other cells at some stage of their lives to establish and maintain infection (Lopez et al., 2011; Oberholzer et al., 2010). Most of the research done in this field has specifically focused on analyzing the communication of the parasites with their host and limited number of studies analyzed the communication between parasites. Although studies of protozoan parasites generally consider them as individual cells in suspension cultures or animal models of infection, it has been shown that by communicating and acting as a group, unicellular organisms have advantages over individual cells (Lopez et al., 2011; Oberholzer et al., 2010). This communication promotes development, survival, access to nutrients, and improves the pathogens’ mechanism of defense against their host (Ofir-Birin et al., 2017). As example, communication between *T. brucei* parasites modulates the quorum sensing-mediated differentiation of into the tsetse fly transmissible short-stumpy developmental form in the mammalian bloodstream (Mony et al., 2014) as well as social motility exhibited by procyclic forms in the insect vector midgut (Oberholzer et al., 2010). Another interesting example about parasite: parasite communication comes from studies in the malaria parasite Plasmodium where it has been shown that parasites adjusted their sex ratio in response to the presence of unrelated genotypes in the parasite population (Pollitt et al., 2011). Although communication among parasites have clear important implications in biology, it has not yet been deeply studied in *T. vaginalis*. Here, we demonstrated that EVs are involved in communication among different *T. vaginalis* parasite strains. Like other eukaryotes, the research of EVs in protozoan parasites has grown recently, and the data point to the involvement of protozoan EVs in cell communication (Szempruch et al., 2016a; Wu et al., 2018). EVs modulate gene expression and affect signaling pathways that might result in developmental changes and modulation of immune response that impact the course of the infection (Nievas et al., 2020; Ofir-Birin et al., 2017). In this sense, EVs isolated from *Trypanosoma cruzi* trypomastigotes lead to parasite spread and survival (Trocoli Torrecilhas et al., 2009). While *Trypanosoma brucei* EVs promoted parasite entrance into host cells (Atyame Nten et al., 2010; Geiger et al., 2010; Vartak and Gemeinhart, 2007), exosomes also have a role in communication between parasites by affecting social motility and migration (Eliaz et al., 2017). Specifically in *T. vaginalis*, EVs have been shown to play a key role in pathogenesis as it has been demonstrated that small EVs have immunomodulatory properties and modulate parasite adherence to host cells (Y R Nievas et al., 2018a; Olmos-Ortiz et al., 2017; Rai and Johnson, 2019; Twu et al., 2013). Although it has been shown that EVs are key players in *T. vaginalis:* host communication, its role in parasite: parasite communication has not been profoundly analyzed yet. In this sense, a previous study showed that preincubation of small EVs isolated from highly adherent strains increased the attachment of a poorly adherent strain (Twu et al., 2013), suggesting that small EVs might be involved in both parasite: parasite and parasite: host communication. In concordance, we found that EVs are responsible, at least in part, of communication between different parasite strains as the incubation of EVs enriched samples from G3 or B7RC2 have an effect in the formation of filopodia and cytonemes of CDC1132 strain. We have previously demonstrated that EVs enriched samples are obtained using this isolation protocol (Salas et al., 2021), however, it is important to have in mind that coprecipitating proteins might be present in the samples due to limitations of the enrichment techniques used. To better understand the molecular pathways involved in the differential response in cytonemes and filopodia formation induced by EVs, the protein content of EVs secreted by different *T. vaginalis* strains was analyzed by mass spectrometry. Surprisingly, the number of proteins detected in the EVs proteome of B7RC2 strain was consistently lesser than the number of proteins detected in G3 or CDC1132. In concordance with this, previous sEVs and MVs proteomes performed in B7RC2 strain identified 215 and 592 proteins, respectively (Yesica R. Nievas et al., 2018; Twu et al., 2013). However, when sEVs proteome of TV79-49c1 strain was analyzed, the number of proteins detected in these samples were 1633 (Rada et al., 2022). These and our data (Yesica R. Nievas et al., 2018; Rada et al., 2022; Twu et al., 2013) suggest that EVs released from different *T. vaginalis* strains are clearly distinct and may package strain specific proteins. The difference in the number and identities of proteins identified in the different EV proteomes is striking and suggesting that may serve as part of a specific sorting pathway for delivery of active molecules to neighboring cells. However, even though the proteins released in the EVs were different, the biological processes identified by GO analysis were highly conserved. This diversity in protein cargo might explain the differential response observed when different EVs are incubated with CDC1132 receiving cells.

Our results demonstrate that communication among strains involve EVs released by different strains that induce the formation of cytonemes and filopodia in recipient cells (Fig. 7). Cell protrusions are extensions of the plasma membrane of individual cells that function in sensing the environment and making initial, dynamic adhesions to extracellular matrix and to other cells (Adams, 2004). Specifically, filopodia are thought to transmit signaling molecules to neighboring cells (Roy et al., 2014; Sanders et al., 2013), increase adhesion (Albuschies and Vogel, 2013; Fierro-González et al., 2013), and serve as a sensing organelle (Wood and Martin, 2002). Cytonemes are specialized types of signaling filopodia that exchange signaling proteins between cells (Casas-Tintó and Portela, 2019). The latest research in mammalian cells proposes filopodia-like cell protrusion as a novel form of intercellular communication. Similarly, Bloodstream African trypanosomes also produce membranous protrusions that originate from the flagellar membrane and make connections with the posterior ends of other trypanosomes (Szempruch et al., 2016b). The authors also showed that these interactions were stable over long distances (>20 μm) and highly dynamic (Szempruch et al., 2016b). In concordance, we describe here that formation of cytonemes in *T. vaginalis* is affected by the presence of a different parasite strain. Additionally, the communication among parasites from different strains affect their attachment to host cells; most likely as a result of the increased formation in filopodia and cytonemes structures. Considering that the occurrence of mixed *T. vaginalis* infections has been reported (10.9% of cases) (Conrad et al., 2012), our findings may have important clinical repercussions as we observed that a poorly adherent parasite strain (G3) adheres more strongly to prostate cells in the presence of a highly adherent strain. Although the relevance of parasite: parasite communication in *T. vaginalis* has not been deeply explored, these results suggest that interaction among isolates with distinct phenotypes might affect the behavior recipient cells probably affecting the outcome of infection. In concordance, growing empirical evidence obtained from patients and/or animal models demonstrates that multiple-strain infections in different human pathogens can change dynamics, disease course, and transmission (Balmer and Tanner, 2011). Multiple-strain infections have been shown unambiguously in 51 human pathogens and are likely to arise in most pathogen species (Balmer and Tanner, 2011), indicating that multiple-strain infections are probably the norm and not the exception. The study of signaling, sensing and cell communication among parasitic organisms will improve our understanding of host-pathogen interactions and disease dynamics, providing a basis for novel control approaches.

**Figure 7:**
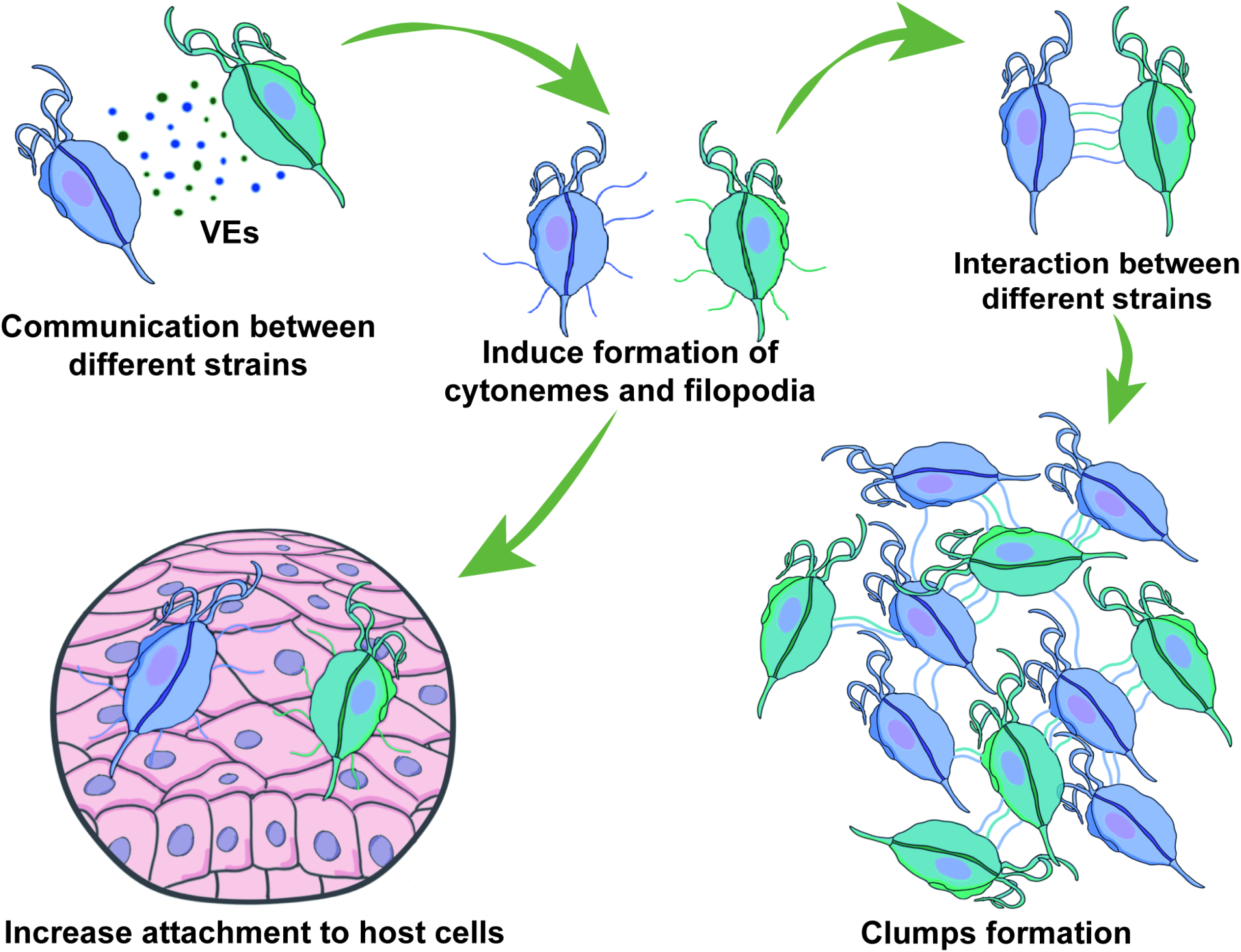
Visual summary of role of EVs and cytonemes in *T. vaginalis* parasite: parasite communication. Different *T. vaginalis* strains release EVs that have specific protein content and affect the formation of filopodia and cytonemes of recipient parasites. The communication among parasites from different strains affect their attachment to host cells; most likely as a result of the increased formation in filopodia and cytonemes structures.

## Supplementary Figures

**Supplementary Figure S1:**
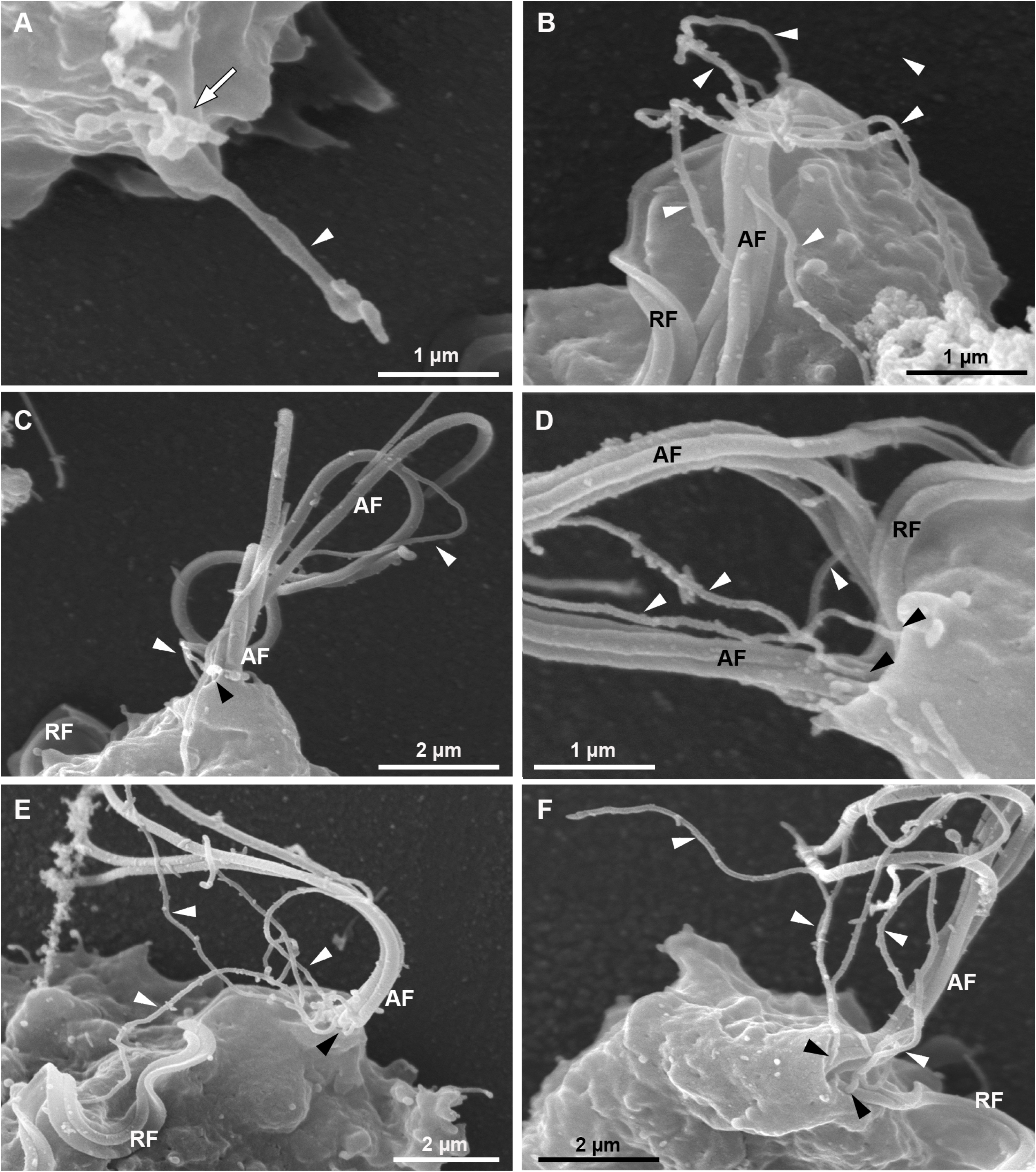
SEM of detailed views of cytonemes protruding from the surface of CDC1132 parasites. Fig. A-D are insets from Fig. 1B. **A**. A tubular projection protruding from the cell body is seen (arrowhead). Notice that the cytoneme is branched in the region where it emerges from the cell surface (arrow). **B-F**. Flagellar cytonemes (white arrowheads). These tubular extensions are originated from the same region where the anterior (AF) and recurrent (RF) flagella emerge (black arrowheads).

**Supplementary Figure S2:**
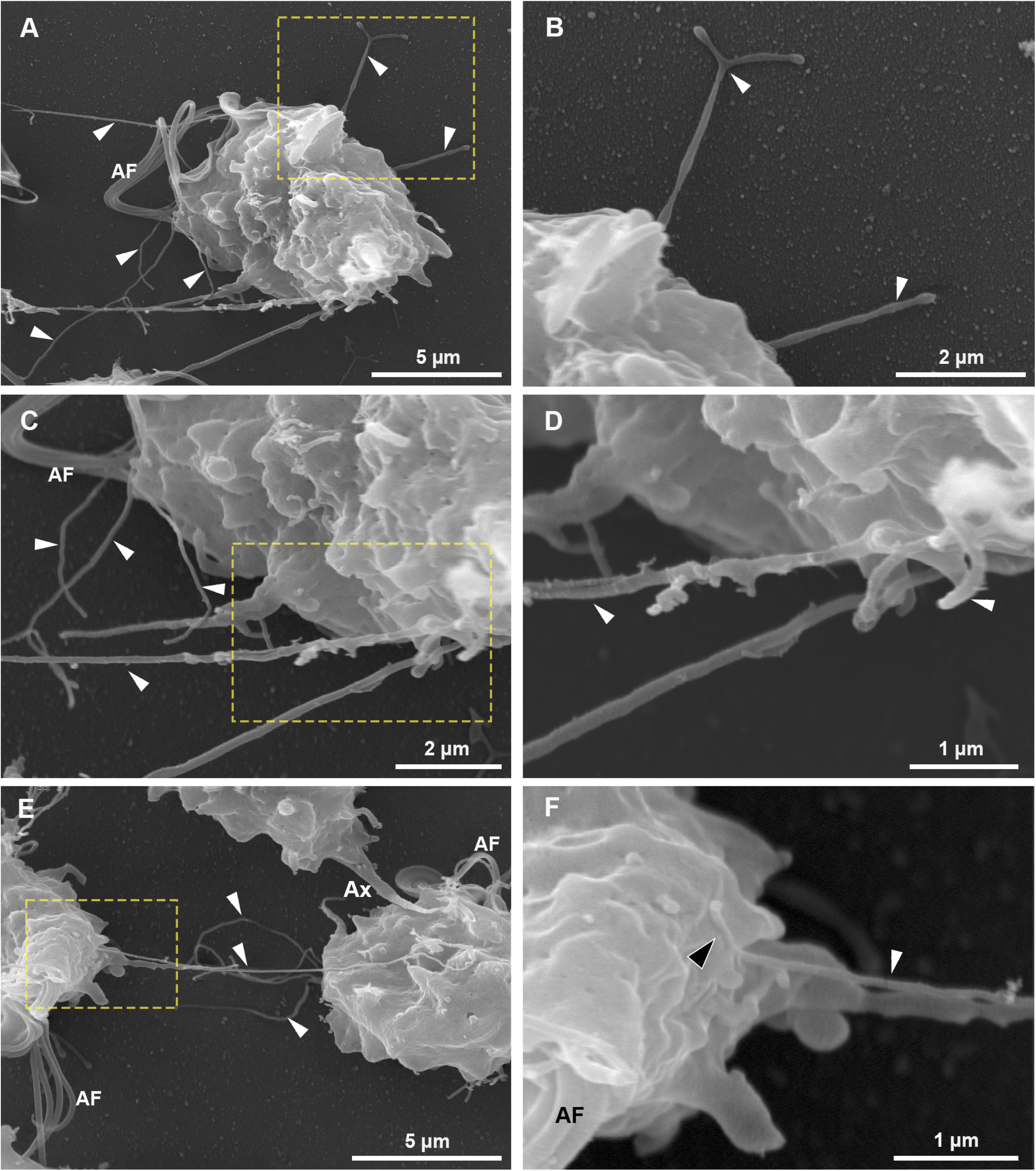
SEM of detailed views and insets from Fig. 2D. **A-D**. Branched and unbranched cytonemes are seen (white arrowheads). **E-F**. A cytoneme connecting two parasites. In **F**, the closed tip of cytoneme is seen in contact with the surface of the adjacent parasite (black arrowheads). AF, anterior flagella; RF, recurrent flagellum.

## Acknowledgements

We thank our colleagues in the lab for helpful discussions. This research was supported with a Grant from the Agencia Nacional de Promoción Científica y Tecnológica (ANPCyT) Grant BID PICT-2019-01671 (NdM) and Arturo Falaschi ICGEB Fellowship (ICGEB). NdM is a researcher from the National Council of Research (CONICET) and UNSAM. NS and MBP are PhD fellow from CONICET. The funders had no role in study design, data collection and analysis, decision to publish, or preparation of the manuscript.

## Author contribution

Natalia de Miguel conceived the study. Experiments were performed by Nehuén Salas, Manuela Blasco-Pedreros, Tuanne dos Santos Melo, Antonio Pereira-Neves, Vanina Maguire and Jihui Sha. The manuscript was drafted by Nehuén Salas and Natalia de Miguel. All authors edited, reviewed and approved the manuscript.

